# Fragility of ER homeostatic regulation underlies haploid instability in human somatic cells

**DOI:** 10.1101/2024.04.05.588249

**Authors:** Sumire Ishida-Ishihara, Kan Yaguchi, Kimino Sato, Koya Yoshizawa, Sena Miura, Guang Yang, Krisztina Veszelyi, Gabor Banhegyi, Eva Margittai, Ryota Uehara

## Abstract

Mammalian somatic cells are generally unstable in the haploid state, resulting in haploid-to-diploid conversion within a short time frame. However, cellular and molecular principles that limit the sustainability of somatic haploidy remain unknown. In this study, we found the haploidy-linked vulnerability to ER stress as a critical cause of haploid intolerance in human somatic cells. Pharmacological induction of ER stress selectively induced apoptosis in haploid cells, facilitating the replacement of haploids by co-existing diploidized cells in a caspase-dependent manner. Biochemical analyses revealed that unfolded protein response (UPR) was activated with similar dynamics between haploids and diploids upon ER stress induction. However, haploids were less efficient in solving proteotoxic status, resulting in a bias toward a proapoptotic mode of UPR signaling. Artificial replenishment of chaperone function or inhibition of a UPR signal transducer ATF6 substantially alleviated the haploidy-linked upregulation of proapoptotic signaling and improved haploid cell retention under ER stress. These data demonstrate that the ER stress-driven haploid instability stems from inefficient proteostatic control that alters the functionality of UPR to cause apoptosis selectively in haploids. Interestingly, haploids suffered a higher level of protein aggregation even in unperturbed conditions, and the long-term stability of the haploid state was significantly improved by alleviating their natural proteotoxicity. Based on these results, we propose that the haploidy-specific vulnerability to ER stress creates a fundamental cause of haploid intolerance in mammalian somatic cells. Our findings provide new insight into the principle that places a stringent restriction on the evolution of animal life cycles.

## Introduction

Mammalian species invariably have a diplontic life cycle, where the multicellular somatic stage is restricted to the diploid state (i.e., each cell possesses two genome copies). Mammalian haploid somatic cells occasionally arising from irregular biological events, such as parthenogenesis or tumorigenesis, cause developmental defects or tissue homeostatic disruption, respectively (1). Mammalian haploid somatic cells are generally unstable and prone to cell division failure that converts them to diploid through whole-genome duplication (1-5). Moreover, haploid cells suffer poorer proliferative potential than their diploidized derivatives (5,6). As a result, haploid cell population is eventually replaced by diploid population during consecutive cell proliferation either in vitro or in vivo (5,7). This haploid intolerance indicates that certain aspects of somatic cell regulations can be maintained only in the diploid state in mammals. However, the molecular bases of the haploid intolerance in mammalian somatic cells are largely unknown.

A previous screening for compounds affecting haploid stability found 3-hydroxy-3-methylglutaryl-coA reductase (HMGCR) inhibitors statins to facilitate the haploid-to-diploid conversion of HAP1 cells (8). Statin-mediated haploid destabilization occurred not through cholesterol depletion, a primary downstream target of statins, but through evoking endoplasmic reticulum (ER) stress (9). However, the pleiotropic effects of statins on broad cellular processes precluded further investigation of the link between ER homeostatic control and haploid instability in human cells. Therefore, molecular and cellular principles potentially destabilizing the haploid state under ER stress are entirely unknown. Clarifying these issues would elucidate key aspects of cell regulations that limit the sustainability of the haploid state in the somatic stage in the mammalian life cycle.

ER stress management is governed by unfolded protein response (UPR) (10). In mammals, UPR consists of three signaling pathway branches mediated by different ER transmembrane proteins, PKR like ER kinase (PERK)/Eukaryotic translation initiation factor 2α kinase 3 (EIF2AK3), Inositol requiring enzyme 1 (IRE1), and Activating transcription factor 6 (ATF6) (11). In non-stressed conditions, these signaling transducers are sequestered in inactive states by direct interaction with an ER chaperone, Binding immunoglobulin protein (BiP)/Glucose-related protein 78 (Grp78)/Heat shock protein family A member 5 (HSPA5) (12-14). Upon the accumulation of unfolded proteins in the ER lumen, pools of ER chaperones, including BiP, are recruited for chaperoning the unfolded proteins, resulting in the release and activation of these UPR transducers.

PERK is activated through dimerization and autophosphorylation. Active PERK then phosphorylates Eukaryotic translation initiation factor 2A (eIF2α) to suppress general mRNA translation and reduce the burden of nascent protein folding (15). Meanwhile, phosphorylated eIF2α selectively increases translation of activating transcription factor 4 (ATF4) (16), which in turn triggers increased transcription of numerous genes associated with ER stress management, including ER chaperones and autophagic factors (17-19). IRE1 is also activated through dimerization and autophosphorylation. Active IRE1 splices X-box-binding protein 1 (XBP1) mRNA, inducing the translation of the active form of XBP1 (20). XBP1 then mediates transcription of numerous genes involved in alleviating ER stress, such as ER chaperones and ER-associated degradation (ERAD) factors for clearance of misfolded protein (21). Activation of ATF6 is mediated through its relocation from ER to the Golgi apparatus, where it is cleaved by Golgi-resident Site-1 and Site-2 proteases (22,23). PERK is also involved in the full activation of ATF6 (24). Then, the cleaved form of ATF6 released to the cytoplasm is translocated into the nucleus to mediate transcription of XBP1, chaperones, and ERAD factors (20,25-28). These reactions mediated by UPR pathways alleviate ER stress and promote cell survival.

However, when ER stress is prolonged or unmanageable, UPR changes its function to drive apoptosis and remove damaged cells. A proapoptotic transcription factor, C/EBP homologous protein (CHOP)/Growth arrest and DNA damage inducible gene 153 (GADD153) is a main mediator of ER stress-induced apoptosis (29-32), which is transcriptionally upregulated mainly by ATF4 and supportively by ATF6 (16,25,33). Therefore, the roles of UPR are highly cell context-dependent, but cellular conditions determining the balance between the pro-survival and proapoptotic function of UPR remain largely unknown (34,35).

In this study, we found that haploid human somatic cells are significantly less efficient in solving ER stress and more prone to ER stress-induced apoptosis than isogenic diploid counterparts. Comparative biochemical analyses revealed that the induction of intermediate levels of ER stress through different stressors activated proapoptotic signaling preferentially in haploids in an ATF6-dependent manner. This ploidy-dependent alteration in UPR functionality drove the rapid expansion of co-existing diploid over haploid cell population, lowering the stability of the haploid state under ER stress. Interestingly, haploid cells suffered a higher level of proteotoxicity than diploids even in unperturbed conditions, alleviation of which by a chemical chaperone significantly resolved the innate instability of the haploid state. These findings indicate that haploidy-linked proneness to ER stress limits the proliferative capacity in the haploid state, creating a fundamental cause of the haploid instability in mammalian somatic cells.

## Results

### ER stress induction aggravates haploid instability through a haploidy-selective cell proliferation suppression

We previously found that statin impaired the stability of the haploid state by evoking ER stress in a human haploid model cell line, HAP1 (9). However, the pleiotropic effects of mevalonate pathway suppression by statin precluded further investigation of a possible relationship between ploidy status and ER homeostatic controls. To address this issue, we tested the effects of more specific ER stress inducer tunicamycin (an inhibitor of N-glycosylation) on the stability of the haploid state in HAP1 cells.

We first treated purified haploid cells with or without tunicamycin and compared the lifetime of the haploid population between the conditions during consecutive passages (Fig. 1A, B, and S1A). In the untreated condition, the initially pure haploid population gradually converted to diploids in successive passages for a few weeks, resulting in a reduction of haploid G1 proportion with the emergence of diploid G2/M proportion in flow cytometric DNA content analysis (Fig. 1A and S1A). Treatment with 50 nM tunicamycin, which allowed cell proliferation with a detectable level of UPR (Fig. S1B; see also Fig. 3), further destabilized the haploid state during the long-term culture (Fig. 1A). Ratio of diploid G2M to haploid G1 population became 4.5 times higher in tunicamycin-treated culture than in control after 20 d (Fig. 1B see also Fig. 6C for more extended culture). Tunicamycin treatment did not cause tetraploidization (Fig. 1A, see also Fig. 6C), demonstrating that it specifically facilitated diploid cell expansion rather than causing general polyploidization. Tunicamycin at a higher concentration than 100 nM completely blocked cell proliferation, precluding the investigation of its effects on long-term haploid stability (Fig. S1B).

**Figure 1:**
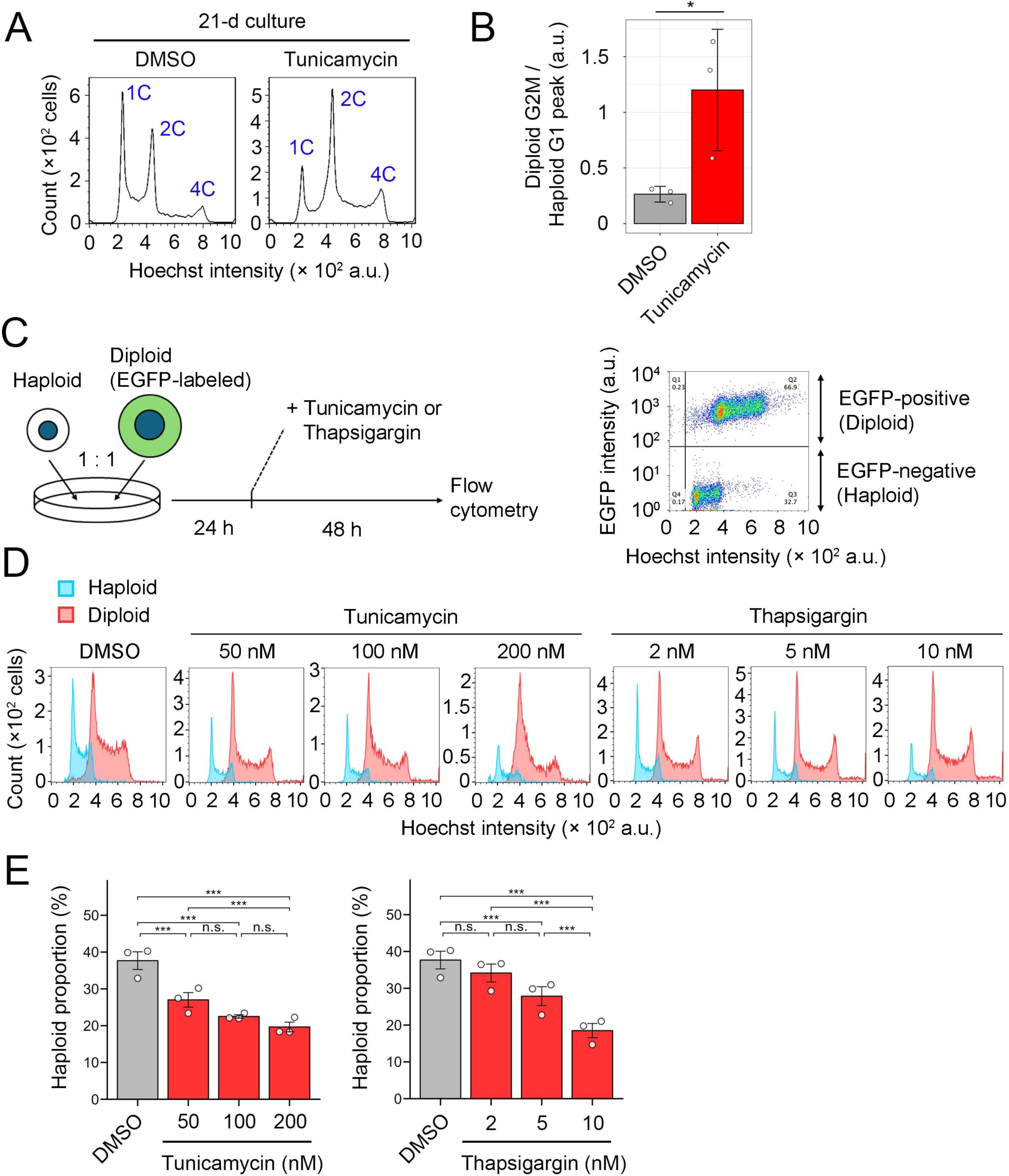
ER stress induction causes destabilization of the haploid state in HAP1. **(A)** Flow cytometric analysis of DNA content in originally haploid HAP1 culture after consecutive passages in the absence or presence of 50 nM tunicamycin for 21 d. DNA was stained by Hoechst 33342. The labels on the plot (1C, 2C, and 4C) are relative DNA amounts (C-value) corresponding to haploid G1, haploid G2M/diploid G1, and diploid G2M, respectively. An example of DNA content analysis in the haploid cell culture before long-term passages (i.e., day 0) is shown in Fig. S1A. **(B)** Ratio of diploid G2M to haploid G1 populations in A. Mean ± SE of 3 independent experiments (sampled at day 20, 21, or 23 of the consecutive culture). The asterisk indicates a statistically significant difference between conditions (\**p* < 0.05, the Brunner-Munzel test). **(C)** Scheme of haploid-diploid co-culture experiment. An example of a dot plot of EGFP intensity against the Hoechst signal in the co-culture flow cytometric analysis is shown on the right. Cell populations originating from haploid or diploid cells were distinguished based on EGFP signal intensity for the analyses in D and E. **(D)** DNA content analysis of co-cultured haploid and diploid cells treated with different concentrations of tunicamycin or thapsigargin for 48 h. Representative data from 3 independent experiments. **(E)** The proportion of haploid cells in the haploid-diploid co-culture. Mean ± SE of 3 independent experiments for each condition. Asterisks indicate statistically significant differences between conditions (n.s.: not significant, \*\*\**p* < 0.001, the Steel-Dwass test). For comparison, identical control data (DMSO) are shown in each graph in E.

Next, we sought to assess the possible deleterious effects of ER stress on the stability of the haploid state in a shorter period. For this, we conducted co-culturing of haploid HAP1 cells with their isogenic diploids labeled with EGFP expression at a 1:1 ratio in the presence of different concentrations of tunicamycin and analyzed changes in haploid proportion after 48-h incubation using flow cytometry (Fig. 1C). Even in untreated control, haploid proportion was reduced to 38% after 48 h, reflecting the less efficient proliferation of haploids than diploids (Fig. 1D and E) (5). Tunicamycin significantly reduced haploid proportion compared to untreated control in a dose-dependent manner (Fig. 1D and E). Importantly, the DNA content of the originally haploid population did not increase by tunicamycin treatment during 48-h incubation (Fig. 1D). This result indicates that ER stress selectively suppresses haploid cell proliferation and helps expansion of diploidized population in originally haploid cell culture, rather than facilitating whole-genome duplication of haploid cells. A significant reduction in the haploid proportion in the co-culture was also observed when treated with another ER stress inducer, thapsigargin (an inhibitor of intracellular Ca^2+^ transport; Fig. 1D and E). Therefore, the haploidy-selective anti-proliferative effects of ER stress inducers were common across different modes of action. As this haploidy-linked sensitization to ER stress was novel, we further addressed the principle underlying this phenomenon.

### Haploidy-linked aggravation of apoptosis underlies the haploidy-selective proliferation suppression under ER stress

We reasoned that ER stress inducers reduced haploid proportion in the haploid-diploid co-culture through selective inhibition of cell cycle progression or induction of cell death. The DNA content analysis showed no difference in cell cycle distribution between haploids and diploids treated with ER stress inducers (Fig. 1D). Therefore, we next conducted annexin V-FITC staining assay to compare the frequency of early apoptotic cells between haploid and diploid HAP1 cell cultures treated with 50 nM tunicamycin for 48 h (Fig. 2A and B). In non-treated control, only a small cell proportion was annexin V-FITC positive in haploid and diploid cell cultures (6.9% and 3.0%, respectively; Fig. 2B), with a slightly higher frequency in haploids. The proportion of annexin V-FITC-positive cells drastically increased by tunicamycin in haploids, while it remained at a low level in diploids (21% and 7.9% in haploids and diploids, respectively; Fig. 2B). Therefore, haploid cells were more prone to apoptosis compared to diploids under low dose ER stress inducer treatment.

**Figure 2:**
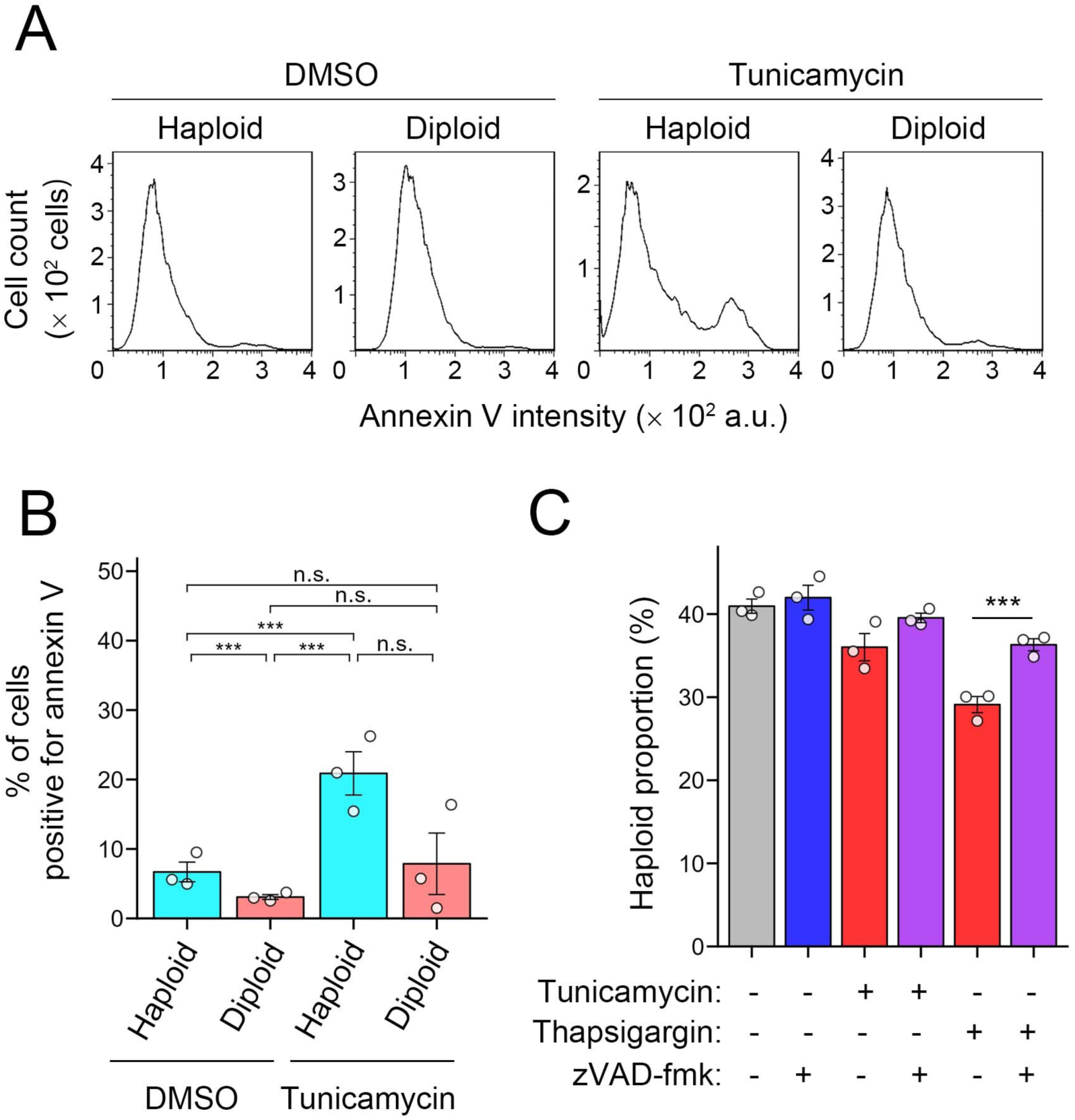
Haploidy-linked aggravation of apoptosis under ER stress. **(A, B)** Flow cytometric analysis of annexin V-FITC staining in haploid or diploid HAP1 cells treated with or without 50 nM tunicamycin for 48 h. Histogram of annexin V-FITC intensity (A) and proportion of annexin V-FITC-positive cells (B). Mean ± SE of 3 independent experiments. Asterisks indicate statistically significant differences among samples (\*\*\**p* < 0.001, the Steel-Dwass test). **(C)** The proportion of haploid cells in co-culture of unlabeled haploid and EGFP-labeled diploid cells treated by tunicamycin or thapsigargin with or without zVAD-fmk for 49 h. Mean ± SE of 3 independent experiments for each condition. Asterisks indicate a statistically significant difference between conditions (\*\*\**p* < 0.001, the Brunner-Munzel test).

To test the causality between the haploidy-liked aggravation of apoptosis and the destabilization of the haploid state under ER stress, we tested the effects of a caspase inhibitor, zVAD-fmk, on haploid proportion in the haploid-diploid co-culture treated with 50 nM tunicamycin or 10 nM thapsigargin (Fig. 2C; see *Experimental procedures*). zVAD-fmk restored the haploid proportion in tunicamycin- or thapsigargin-treated co-cultures, demonstrating that apoptosis blockage alleviates the ploidy-dependent bias of cell proliferation under ER stress (Fig. 2C). These results suggest that the haploidy-linked aggravation of apoptosis is a primary reason of the haploid destabilization under ER stress.

### UPR is biased toward a proapoptotic mode in haploids under ER stress

To specify the molecular basis of the haploidy-linked aggravation of apoptosis upon ER stress, we compared the expression and post-translational modifications of different components of UPR between haploids and diploids treated with different concentrations of tunicamycin for 24 h using immunoblotting (Fig. 3A). Tunicamycin treatment induced UPR events, including increased expression of ER chaperones, BiP and GRP94/HSP90B1/endoplasmin, phosphorylation of PERK with upregulation of its downstream factors, ATF4, and cleavage of ATF6 in a dose-dependent manner both in haploids and diploids without detectable difference between them (Fig. 3A and B). IRE1 remained unchanged in all conditions, both in haploids and diploids (Fig. 3A). Therefore, many UPR components were equivalently reactive to ER stress induction between the haploid and diploid states.

**Figure 3:**
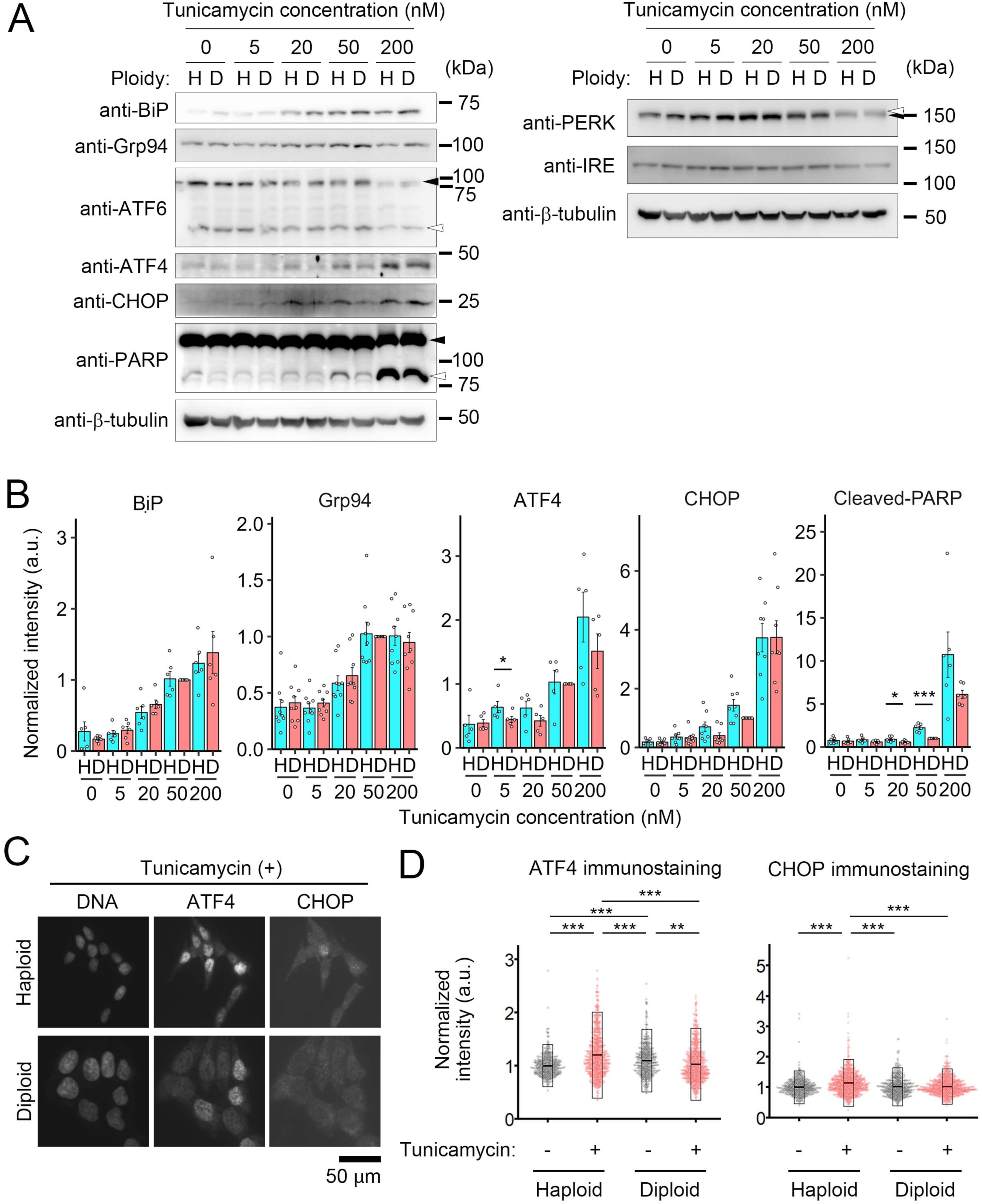
Haploidy-linked upregulation of CHOP expression and PARP cleavage under ER stress. **(A)** Immunoblotting of UPR components and proapoptotic factors in haploid or diploid HAP1 cells treated with different concentrations of tunicamycin for 24 h. Uncleaved or cleaved molecular species are indicated by closed or open arrowheads, respectively, for ATF6 and PARP. Unshifted or shifted PERK bands are indicated by a closed or open arrowhead, respectively. β-tubulin was detected as a loading control. Representative results from ≥3 independent experiments. **(B)** Quantification of the relative intensity of BiP, Grp94, ATF4, CHOP, or cleaved PARP. Mean ± SE of ≥5 independent experiments. Asterisks indicate statistically significant differences among samples (\**p* < 0.05, \*\*\**p* < 0.001, the Brunner-Munzel test). **(C)** Immunofluorescence microscopy of ATF4 and CHOP in haploid or diploid HAP1 cells treated with 50 nM tunicamycin for 24 h. DNA was stained by DAPI. **(D)** Quantification of ATF4 or CHOP signal at nuclei in C. Mean ± SE of ≥602 cells from 2 independent experiments. Asterisks indicate statistically significant differences among samples (\*\**p* < 0.01, \*\*\**p* < 0.001, the Tukey test).

While we did not detect ploidy-dependent differences in the UPR components mentioned above, we found a trend of higher expression of ATF4 or CHOP in haploids compared to diploids treated with different concentrations of tunicamycin (Fig. 3B). To further confirm this trend at single cell level, we also conducted co-immunostaining of ATF4 and CHOP in tunicamycin-treated haploids and diploids (Fig. 3C). Nuclear levels of ATF4 and CHOP staining positively correlated both in haploids and diploids (Fig. S2A) and were significantly higher in haploids than in diploids after 24-h treatment with tunicamycin (Fig. 3D). Therefore, upregulation of ATF4-CHOP axis was more drastic in haploids than in diploid under modest ER stress. Since increased CHOP expression implied upregulation of proapoptotic signaling, we also compared the extent of PARP cleavage, a common consequence of proapoptotic signal activation under unsolved ER stress (36). Consistent with the haploidy-biased CHOP upregulation, haploids manifested a significantly higher magnitude of PARP cleavage than diploids in a low-concentration range of tunicamycin (Fig. 3B). Similar trends of the haploidy-linked aggravation of the proapoptotic signaling was also observed upon treatment with different concentrations of thapsigargin (Fig. S2B and C). Therefore, haploid cells were more prone to proapoptotic status than diploids upon UPR activation under moderate ER stress, consistent with the result in the annexin V binding assay (Fig. 2A and B).

To obtain insight into the dynamics of the aggravation of proapoptotic signaling in haploid cells under ER stress, we next analyzed time course of expressions of BiP, CHOP, and PARP cleavage during 24-h tunicamycin treatment in haploids and diploids (Fig. 4A). After 50 nM tunicamycin administration, the expression of BiP monotonically increased with similar kinetics between haploid and diploid cells. On the other hand, the expression of CHOP peaked around 9-12 h after tunicamycin administration and gradually decreased afterward until 24 h, both in haploids and diploids (Fig. 4A and B). However, during 12-24 h, haploid cells kept a higher level of CHOP expression than diploids (Fig. 4B). Consistent with the prolonged CHOP expression, haploid cells manifested a substantially higher level of cleaved PARP than diploids during 12-24 h after tunicamycin administration (Fig. 4A and B). Therefore, while UPR commenced and progressed with similar dynamics in haploids and diploids, haploids were less efficient in recovering from the acute ER stress status.

**Figure 4:**
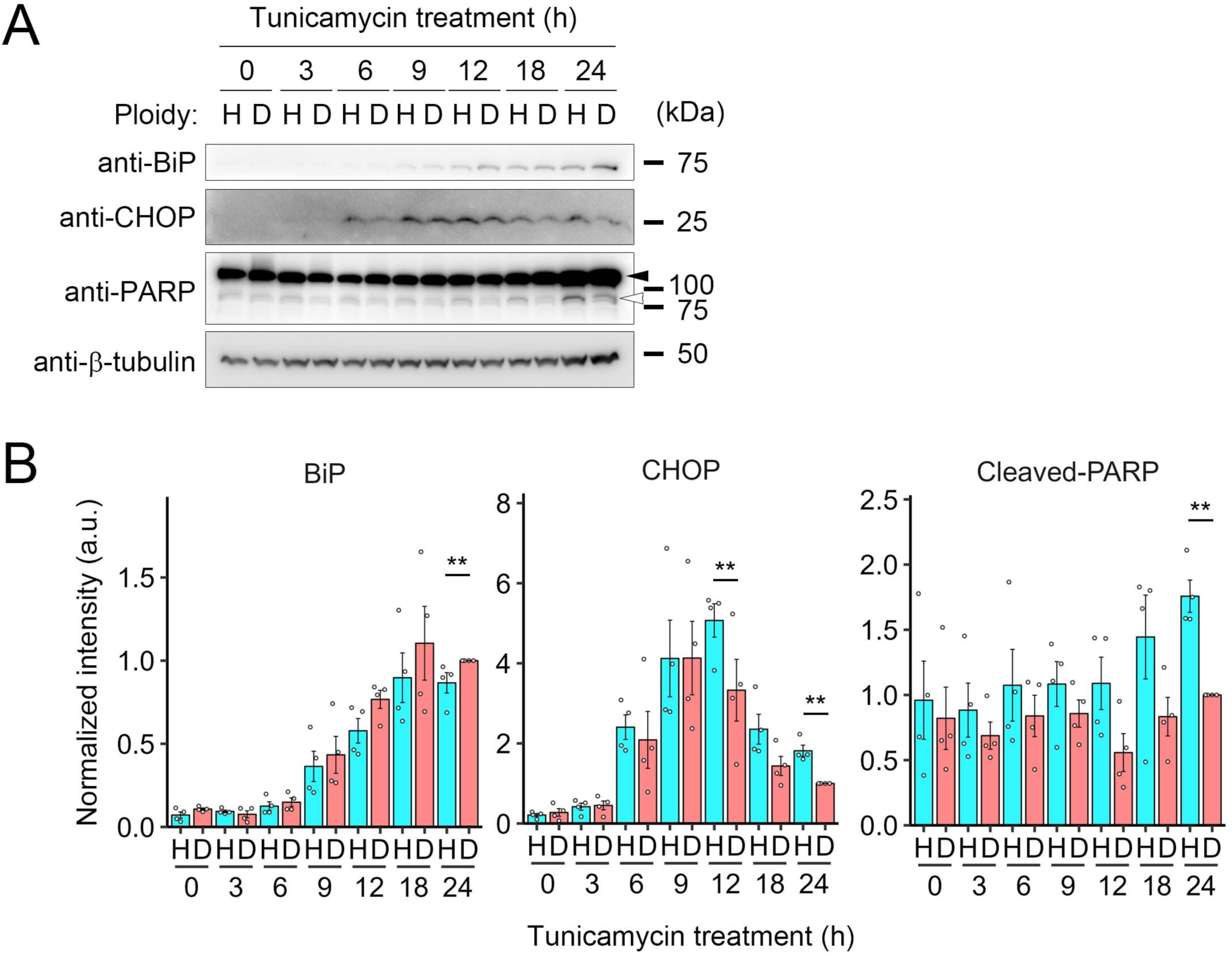
Haploids are less efficient in resolving CHOP signaling upon acute ER stress. **(A)** Immunoblotting of BiP, CHOP, and PARP in haploid or diploid HAP1 cells treated with 50 nM tunicamycin for the indicated duration. The closed or open arrowhead indicates uncleaved or cleaved PARP, respectively. β-tubulin was detected as a loading control. Representative results from 4 independent experiments. **(B)** Quantification of the relative intensity of BiP, CHOP, or cleaved PARP. Mean ± SE of 4 independent experiments. Asterisks indicate statistically significant differences among samples (\*\**p* < 0.01, the Brunner-Munzel test).

### Haploid cells suffer a higher level of protein aggregation either in the presence or absence of ER stressors

The above findings raised a possibility that haploid cells were less efficient in resolving ER stress, hence suffering prolonged stress status to induce proapoptotic signaling. To test this idea, we next assessed the extent of proteotoxicity in ER stress-induced haploids and diploids by post-fixation cell staining using Proteostat dye that specifically labeled misfolded and aggregated proteins (37). We first tested the sensitivity of the dye by comparing its intensity in haploid and diploid cells treated with or without 10 μM MG132 (Fig. 5A and B), a proteasome inhibitor expected to induce drastic protein aggregation (37). Interestingly, Proteostat staining was significantly more intense in haploids than in diploids in the non-perturbed condition (Fig. 5B). MG132 treatment drastically increased Proteostat intensity both in haploid and diploid cells, confirming the protein misfolding detectability by the assay (Fig. 5B). The effect of MG132 was more drastic in haploids, resulting in significantly stronger Proteostat intensity in haploids than diploids in the presence of MG132 (Fig. 5B).

**Figure 5:**
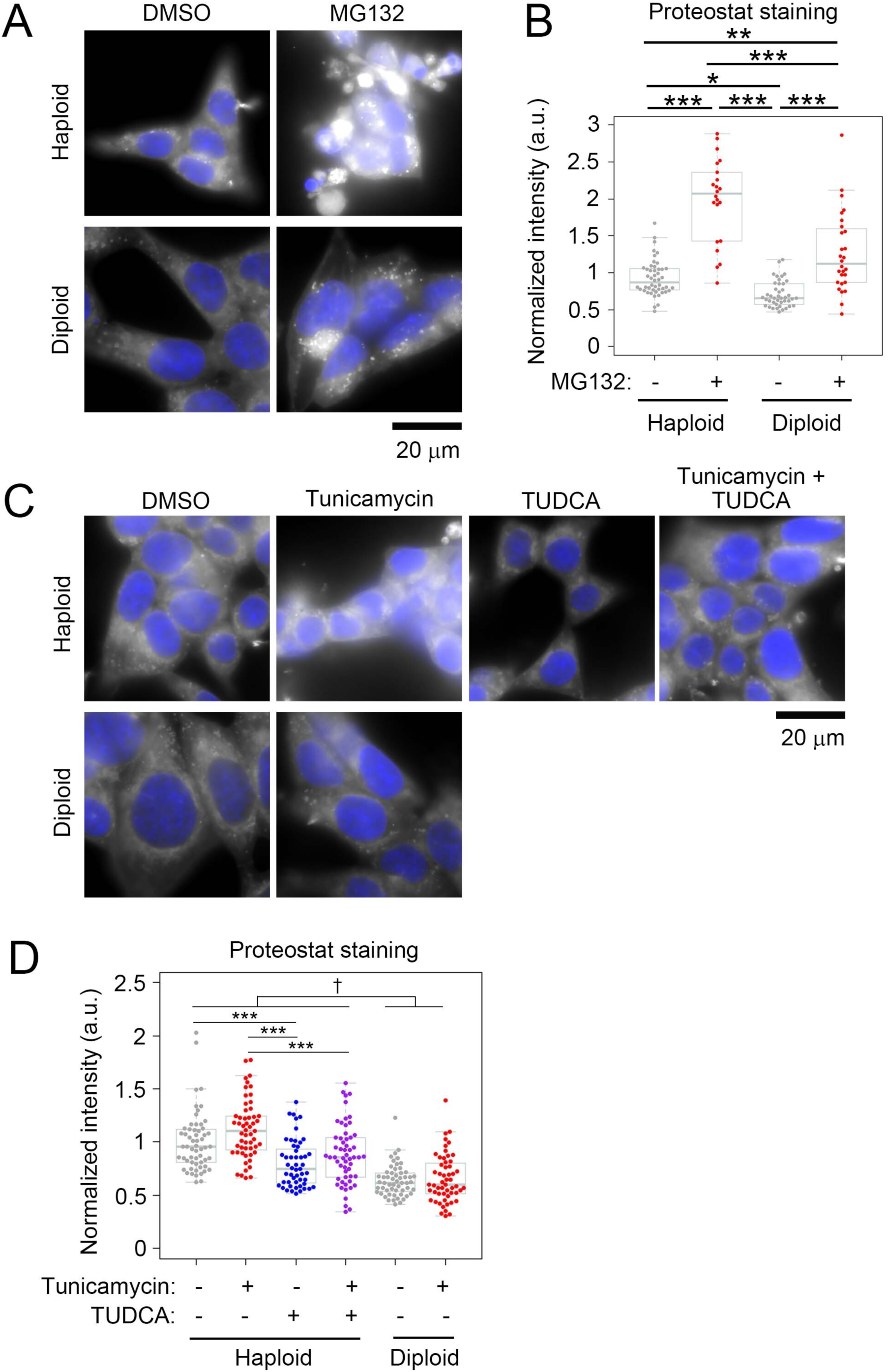
Haploid cells suffer a higher level of protein aggregation. **(A, C)** Fluorescence microscopy of Proteostat dye staining in haploid or diploid HAP1 cells treated with or without 10 μM MG132 (A) or 50 nM tunicamycin in the presence or absence of 2.5 mM TUDCA (C). MG132 was treated for 12 h. Tunicamycin was treated for 24 h, and TUDCA was added at 12 h after the administration of tunicamycin. DNA was stained by Hoechst 33342 (shown in blue). **(B, D)** Quantification of cytoplasmic Proteostat signal in A or C. At least 22 or 51 cells (for B or D, respectively) from 2 independent experiments were analyzed for each condition. Asterisks and a dagger indicate statistically significant differences among samples (\**p* < 0.05, \*\**p* < 0.01, \*\*\**p* < 0.001, †: p<0.001 for each pair between these two groups except between TUDCA-treated haploid vs non-treated diploid (p<0.01) or TUDCA-treated haploid vs tunicamycin-treated diploid (p<0.05), the Tukey test).

Next, we compared Proteostat intensity between haploid and diploid cells treated with or without 50 nM tunicamycin (Fig. 5C and D), which specifically activated proapoptotic signaling in haploids (Fig. 3). In diploids, tunicamycin treatment did not significantly change Proteostat intensity with a slight reduction in the average value from non-treated control (Fig. 5D). This result suggests that UPR upregulation observed in this condition (Fig. 3 and 4) sufficiently resolves proteotoxic state upon tunicamycin treatment in the diploid state. In contrast, Proteostat intensity modestly increased upon tunicamycin treatment in haploids (Fig. 5D). Therefore, haploids were less efficient in solving protein misfolding and aggregation under ER stress. Moreover, the finding that haploids manifested higher Proteostat intensity even in unperturbed conditions demonstrated the elevation of basal proteotoxicity in the haploid state compared to diploids.

### A chemical chaperone restores the stability of haploid cells treated either in the presence or absence of tunicamycin

We next addressed the causality between the poorer ability to solve protein aggregation and proneness to proapoptotic signaling in haploid cells under ER stress. For this, we attempted to restore protein folding capacity in haploid cells by treating them with a chemical chaperone TUDCA and tested its effects on proapoptotic signaling in haploid cells. We introduced TUDCA in haploid cell culture at 12 h after administering 50 nM tunicamycin and tested the time course of CHOP expression and PARP cleavage (Fig. 6A and B). Prior to this analysis, we confirmed that the introduction of TUDCA in this schedule significantly reduced Proteostat signal in haploids either in the presence or absence of tunicamycin (Fig. 5D), showing that TUDCA was indeed effective in facilitating the resolution of either basal or artificially induced proteotoxic stress in the haploid state. Consistent with the results in Fig. 3, CHOP expression peaked at 12 h post-tunicamycin administration. However, TUDCA treatment from that time substantially suppressed CHOP expression at 24 h (Fig. 6B). Importantly, TUDCA also reduced PARP cleavage at 24 h in haploids to the level equivalent to diploids (Fig. 6B). Therefore, even when initial CHOP expression was fully induced, the later replenishment of chaperone function sufficiently canceled the haploidy-linked aggravation of proapoptotic signaling. This data indicates that the lower efficiency in ER stress alleviation is the main cause of proapoptotic signal activation in haploid cells.

**Figure 6:**
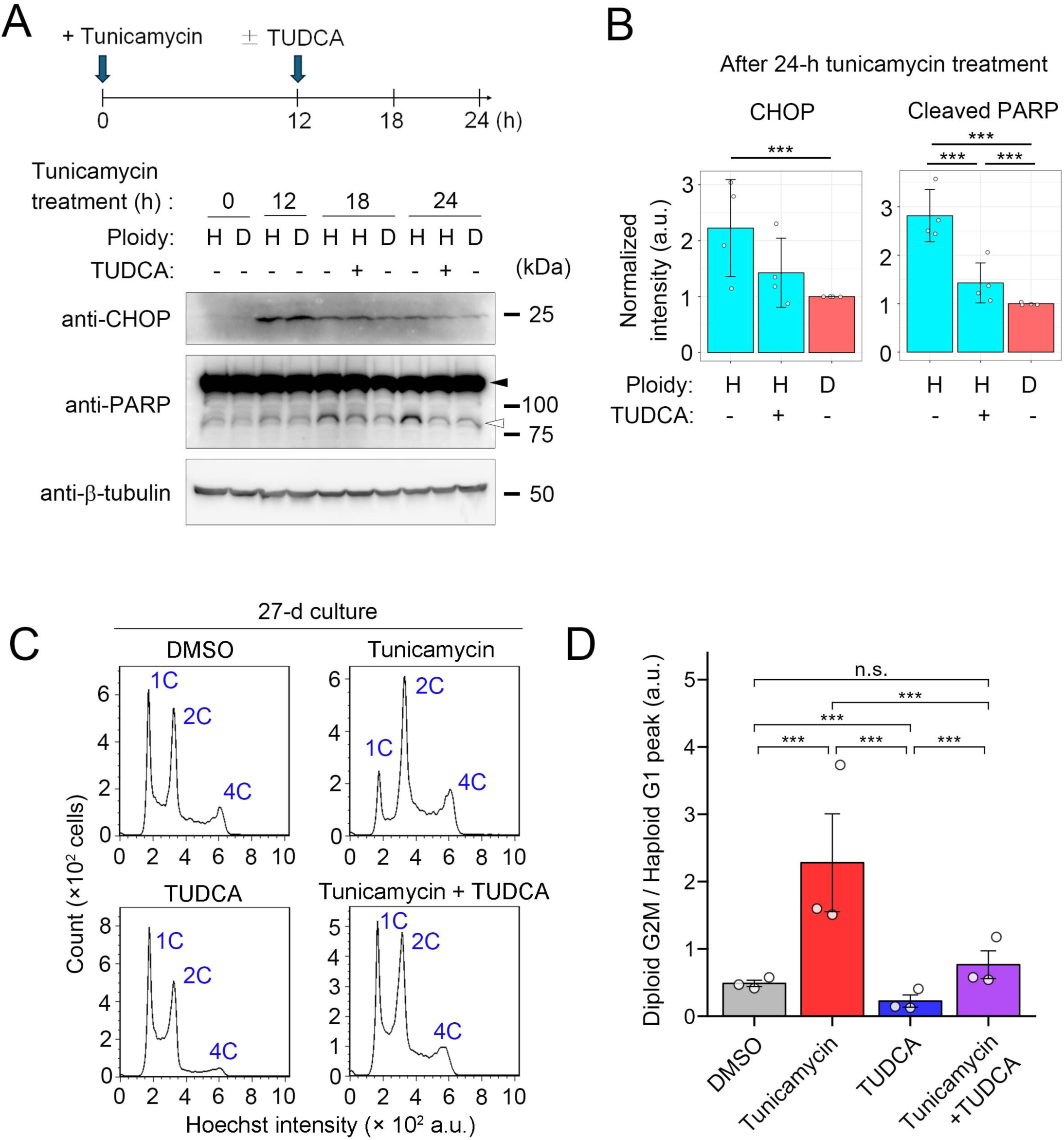
TUDCA stabilizes the haploid state under ER stress. **(A)** Immunoblotting of CHOP and PARP in haploid or diploid HAP1 cells treated with 50 nM tunicamycin for the indicated duration. TUDCA was administrated to haploid cell culture at 12 h after the introduction of tunicamycin, as depicted in the scheme on top. The closed or open arrowhead indicates uncleaved or cleaved PARP, respectively. β-tubulin was detected as a loading control. Representative results from 4 independent experiments. **(B)** Quantification of the relative intensity of CHOP, or cleaved PARP at the 24-h time point in A. Mean ± SE of 4 independent experiments. Asterisks indicate statistically significant differences among samples (\*\*\**p* < 0.001, the Steel-Dwass test). **(C)** Flow cytometric analysis of DNA content in originally haploid HAP1 culture after consecutive passages in the absence or presence of 50 nM tunicamycin or TUDCA for 27 d. DNA was stained by Hoechst 33342. An example of DNA content analysis in the haploid cell culture before long-term passages (i.e., day 0) is shown in Fig. S1A. **(D)** Ratio of diploid G2M to haploid G1 populations in C. Mean ± SE of 3 independent experiments (sampled at day 27 of the consecutive culture). Asterisks indicate statistically significant differences from the control (\*\*\**p* < 0.001, the Steel-Dwass test).

Next, we tested whether the replenishment of the chaperone function by TUDCA restored the stability of the haploid state in tunicamycin-treated long-term cell culture (Fig. 6C and D). Consistent with the previous data (Fig. 1A), tunicamycin treatment accelerated the haploid-to-diploid conversion during 27-d culture. However, the acceleration of the haploid-to-diploid conversion was canceled by co-treating TUDCA with tunicamycin (Fig. 6C and D). Moreover, we found that a single treatment of TUDCA significantly stabilized the haploid state during the long-term culture when compared to non-treated control (Fig. 6D). Considering that haploids suffered severe basal proteotoxic stress (Fig. 5B and D), this result indicates that not only artificially induced ER stress but also basal ER stress in unperturbed haploid cells contributes to the instability of the haploid state in mammalian somatic cells.

### ATF6 contributes to the ploidy-dependent cell proliferation bias

Our data above indicate that the poorer ability of haploids to solve proteotoxicity accounts for the shift of the UPR functionality toward proapoptotic mode and selective cell proliferation suppression in haploids. We reasoned that if particular UPR branches were involved in this process, their suppression would differentially affect the survival and proliferation of haploids and diploids. To test this idea, we analyzed the effects of inhibition of each UPR sensor by GSK2656157 (a PERK inhibitor), 4μ8C (an IRE1 inhibitor), or Ceapin-A7 (an ATF6 inhibitor) on haploid proportion in the haploid-diploid co-culture after 48-h incubation in the presence or absence of 50 nM tunicamycin (Fig. 7A, B, S3A, and B). When singly treated in the co-culture, GSK2656157 or 4μ8C reduced or did not change haploid proportion from non-treated control, respectively (Fig. S3A). On the other hand, Ceapin-A7 significantly increased haploid proportion compared to non-treated control (Fig. 7A). Ceapin-A7 also improved retention of haploid population significantly in the tunicamycin-treated co-culture to the level equivalent to non-treated control (Fig. 7B), while such an effect was not observed with either GSK2656157 or 4μ8C (Fig. S3B).

**Figure 7:**
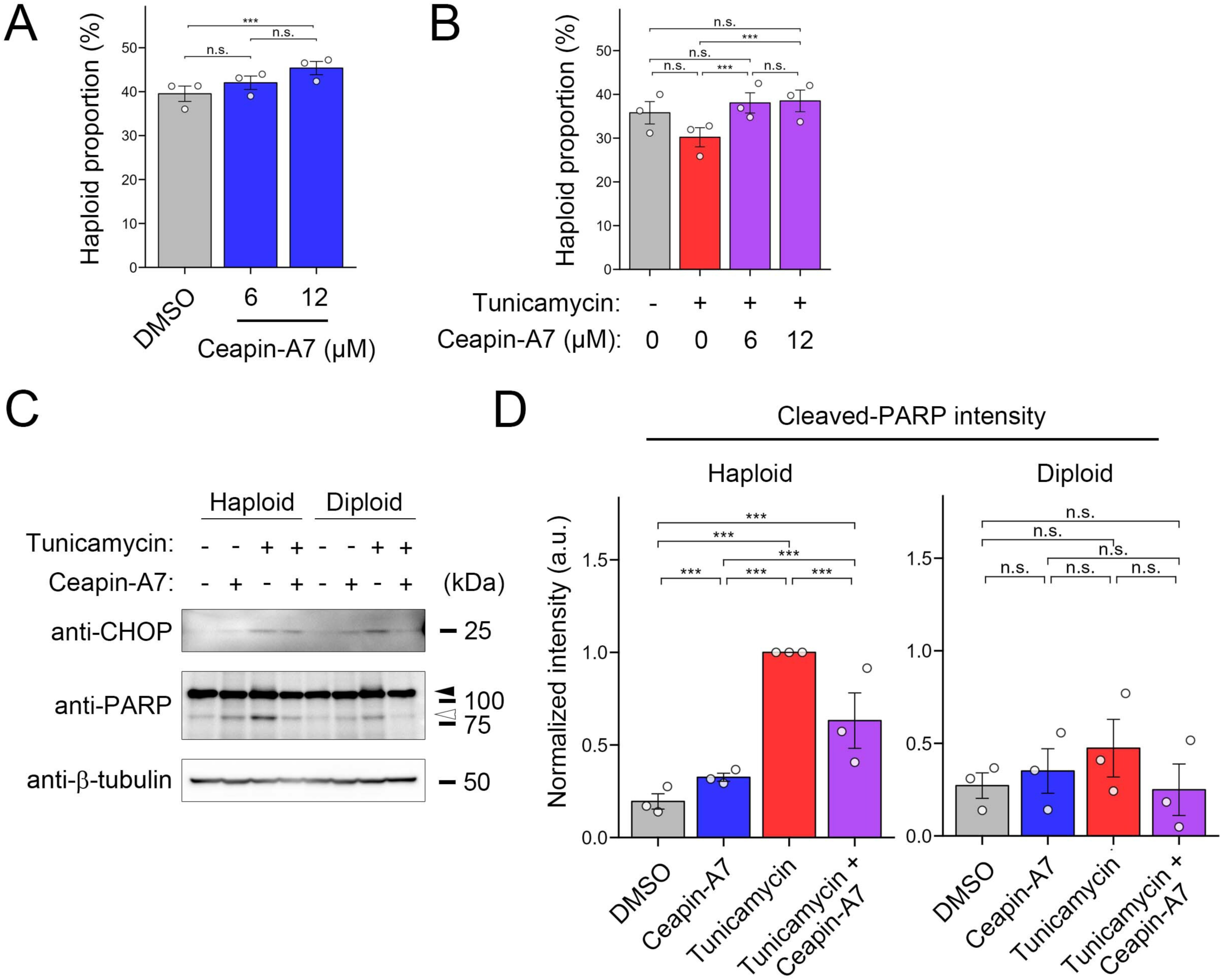
ATF6 is involved in haploidy-selective cell proliferation suppression under ER stress. **(A, B)** The proportion of haploid cells in the haploid-diploid co-culture treated with Ceapin-A7 (A) or co-treated with Ceapin-A7 and tunicamycin (B). Mean ± SE of 3 independent experiments for each condition. Asterisks indicate statistically significant differences between conditions (\*\*\**p* < 0.001, the Steel-Dwass test). **(C)** Immunoblotting of CHOP and PARP in haploid or diploid HAP1 cells treated with 50 nM tunicamycin with or without 12 μM Ceapin-A7 for 24 h. The closed or open arrowhead indicates uncleaved or cleaved PARP, respectively. β-tubulin was detected as a loading control. Representative results from 3 independent experiments. **(D)** Quantification of the relative intensity of cleaved PARP in C. Mean ± SE of 3 independent experiments. Asterisks indicate statistically significant differences among samples (\*\*\**p* < 0.001, the Steel-Dwass test).

ATF6 contributes to pro-survival or proapoptotic reactions under ER stress (38-42). Therefore, inhibition of ATF6 could have canceled ploidy-dependent cell proliferation bias either by suppressing pro-survival activities in diploids or proapoptotic activities in haploids. To distinguish these two possibilities, we tested the effects of Ceapin-A7 on proapoptotic signaling status in haploids and diploids in the presence or absence of tunicamycin (Fig. 7C, D, and S3C). Ceapin A7 treatment did not significantly change CHOP expression level either in the presence or absence of tunicamycin in haploids and diploids (Fig. S3C). In contrast, Ceapin-A7 significantly suppressed PARP cleavage in the presence of tunicamycin in haploid cells (Fig. 7D). Ceapin-A7 also suppressed tunicamycin-dependent PARP cleavage in diploids, although in a much more modest manner compared to haploids (Fig. 7D). These data indicate that, among UPR branches, at least ATF6 is involved in proapoptotic signaling upregulation, contributing to the ploidy-dependent cell proliferation bias.

## Discussion

### Destabilization of the haploid state under ER stress

The instability of the haploid state with gradual haploid-to-diploid conversion is commonly observed in mammalian somatic cells. We previously found two contributors to haploid-to-diploid conversion: spontaneous whole-genome duplication via chromosome missegregation and pooler proliferation of haploid cells compared to diploidized ones (5,43). However, the causes of the poorer proliferation of haploid cells remained largely unknown. In this study, we specified the difference in ER stress tolerance between haploid and diploid cells, which would provide insight into the principle of the ploidy-linked difference in cell proliferation efficiency and the unstable nature of the haploid state in mammalian somatic cells.

Artificial induction of ER stress drastically facilitated haploid-to-diploid conversion in long-term culture experiments. Our results from the haploid-diploid co-culture experiments suggest that destabilization of the haploid state upon ER stress induction is mainly attributed to the haploidy-selective suppression of cell proliferation (Fig. 1). We also found that haploids underwent apoptosis significantly more frequently than diploids under ER stress (Fig. 2), while there was no detectable difference in the effects of ER stress inducers on cell cycle distributions between haploids and diploids (Fig. 1). Moreover, artificial circumvention of apoptosis restored relative proliferative ability of haploids under ER stress (Fig. 2). These findings support the idea that the haploidy-linked aggravation of apoptosis is a primary cause of the selective removal of haploid population that leads to the destabilization of the haploid state under ER stress.

Upon the administration of ER stress inducers, haploid cells activated UPR with similar dynamics as diploids (Fig. 3 and 4). However, haploid cells tended to manifest sustained CHOP expression with higher PARP cleavage levels than diploids. These findings suggest that the UPR mechanism is more biased toward proapoptotic mode in haploid cells than diploids. Interestingly, haploid cells manifested significantly stronger Proteostat staining signals than diploids after the administration of ER stress inducers (Fig. 5), indicating that haploids are less efficient in solving proteotoxic status under ER stress. Since replenishment of chaperone function by TUDCA reduced proapoptotic signaling in tunicamycin-treated haploids to diploid level (Fig. 6), the haploidy-linked bias toward proapoptotic mode of UPR would be mainly attributed to the prolonged ER stress due to the inefficient proteostatic control in haploids.

Among UPR branches, inhibition of ATF6 specifically restored the relative proliferation of haploid cells with alleviation of proapoptotic signaling under artificial ER stress (judged by PARP cleavage; Fig. 7). These data suggest that at least the ATF6 branch is responsible for the haploidy-linked change in the UPR functionality from pro-survival to proapoptotic mode. ATF6 contributes to either pro-survival or proapoptotic signaling depending on the biological contexts and types of stress (38-42). In the context of proapoptotic signaling, while ATF4 mainly mediates upregulation of CHOP expression, ATF6 functions synergistically with ATF4 to tune the CHOP response (33,39). We found that ATF4 and CHOP were more drastically upregulated with positive correlation in haploids under ER stress (Fig. 3). Therefore, an intriguing possibility is that haploidy-linked upregulation of ATF4-CHOP axis, possibly due to the unsolved proteotoxicity, makes the functionality of the ATF6 branch more biased toward proapoptotic in the haploid state. Further study in the future, including comprehensive gene regulatory analyses using transcriptome information, would be required to understand how ATF6-dependent signaling is altered to sensitize the proapoptotic pathway in haploid cells.

### Potential contributions of basal ER stress to haploid instability

It is important to note that haploid cells manifested a higher level of protein aggregation than diploids, even in an unperturbed condition (Fig. 5). We found that TUDCA substantially alleviated the basal protein aggregation and improved the long-term stability of the haploid state compared to the non-treated condition (Fig. 6). These findings indicate an interesting possibility that haploid cells are prone to naturally occurring ER stress that limits the proliferative capacity in the haploid state, creating a fundamental cause of the haploid instability in mammalian somatic cells. Related to this idea, a previous study has reported that deletion of HAC1 gene (a yeast orthologue of XBP1) results in the destabilization of an otherwise stable haploid state in budding yeasts (44), indicating that ER homeostasis is a crucial determinant of sustainability of somatic haploidy in broad species.

Since the basal level of ER stress-driven proapoptotic signaling was too low to be reliably detected by currently available methods, it was impossible to compare it between haploids and diploids accurately. For the same reason, we could not judge whether ATF6 inhibition impacts basal proapoptotic signaling in unperturbed haploid cells in our current study (Fig. 7D and S3C). Therefore, while ATF6 inhibition improved the relative cell proliferation capacity of haploids even in the absence of ER stress inducer (Fig. 7A), it remains unknown whether the ATF6 branch constantly operates as a mechanism that selectively induces apoptosis and limits the proliferative capacity of haploid cells. We expect that molecular mechanisms and physiological effects of the basal ER stress-driven haploid cell suppression may be better addressed in tissue environments with a higher level of basal ER stresses. It would be particularly intriguing to address in the future how the ER stresses that take place in association with developmental or tumorigenic processes affect cellular behavior and tissue functionality in haploid embryogenesis or haploid cancer progression.

### Why are haploid cells less resistant to ER stress?

The reason for the proneness of haploid cells to ER stress and proteotoxicity remains unclear. A possible explanation may be the smaller cell size of haploids. In our previous estimation, haploid HAP1 cells were about half in volume compared to diploids (5). This theoretically results in about a 1.25-time increase in cell surface-to-volume ratio in haploids compared to diploids, potentially increasing the burden of membrane or secreted protein production on haploid cells. Interestingly, previous studies have reported haploidy-linked alterations in the transcription of genes encoding glycoproteins (7,45). The potentially higher demand for ER productivity may cause more frequent protein misfolding in the basal cell state and lead to severe damage upon perturbations of ER functionality in haploid cells. Alternatively, cell size reduction may limit the capacity of ER to manage stresses by limiting its size through organelle size scaling mechanisms (46). Upon UPR activation under ER stress, cells increase their capacity to manage stresses by expanding the ER lumen in addition to enhancing its chaperone activities (47-50). Drastic cell size reduction may limit space for ER to expand upon ER stress and thus decrease the capacity to resolve protein misfolding in haploid cells. A comparative ultrastructural analysis of ER in the presence or absence of ER stress in haploid and diploid cells would provide important insight into this issue in the future.

Besides cell size reduction, the potential insufficiency in gene dosage of critical stress-managing factors may limit the tolerability of haploid cells to ER stress. Consistent with this idea, dosage reduction or increase in specific genes have been demonstrated to alter cellular response to ER stress (51,52). Future studies on the possible influence of cell size or gene dosage would also provide mechanistic insights into the ATF6-dependent aggravation of cell proliferation suppression in haploid cells.

The instability of the haploid state in the multicellular stage in mammals is a fundamental problem in the evolution of animal life cycles. At the same time, this is an important limitation of the application of mammalian haploid cell resources for genetics and bioengineering (53). Our new findings would provide a clue to understanding the principle of haploid intolerance in animals and modulate the stability of mammalian somatic haploidy to increase its utility in life science research fields.

### Experimental procedures

#### Cell culture

Haploid cells (RRID: CVCL_Y019; from Haplogen GmbH) (54) were cultured in Iscove’s Modified Dulbecco’s Medium (IMDM; Wako Pure Chemical Industries) supplemented with 10% fetal bovine serum (FBS) and 1× antibiotic-antimycotic solution (AA; Sigma-Aldrich). Maintenance of haploid HAP1 cell populations using size-based cell sorting and establishment of diploid HAP1 cells were conducted as described previously (5).

For long-term cell passage experiments, freshly purified haploid HAP1 cells were cultured in the presence or absence of compounds described elsewhere. Cells were typically passaged every 2 d with replenishment of the compounds until they were subjected to flow cytometric analyses.

#### Compounds and antibodies

Compounds were purchased from the distributors as follows. Tunicamycin: Cell Signaling Technology or Cayman Chemical. Thapsigargin: Wako. TUDCA: Tokyo Chemical Industry Co., Ltd. GSK2656157, 4μ8C, and Ceapin-A7: Sigma-Aldrich. Z-Val-Ala-Asp(OMe)-CH_2_F (zVAD-fmk): Peptide Institute, Inc. Antibodies used in this study are listed in Table S1.

#### Flow cytometry and cell sorting

Flow cytometric analyses and cell sorting were performed using a JSAN desktop cell sorter (Bay Bioscience). For DNA content analyses, cells were trypsinized using 0.05% trypsin-EDTA solution (Wako) and stained with 10 μg/ml Hoechst 33342 (Dojindo) for 15 min at 37°C. For Annexin V assays, cells were suspended by treating with Accutase® solution (PromoCell) and stained with Hoechst and 1 μg/ml Annexin V-FITC (Medical and Biological Laboratories) in binding buffer (10 mM HEPES, pH 7.4, 140 mM NaCl, 2.5 mM CaCl_2_) for 15 min at 37°C. The stained cells were filtered through a cell strainer and applied to the cell sorter. Data analyses were conducted using FlowJo software (BD Sciences).

#### Co-culture experiment

Non-labeled haploid and EGFP-expressing diploid cells (5 × 10^4^ cells/ml each) were mixed in a 1:1 ratio, 2 ml seeded on 6-well plates coated with collagen type I (Corning). After 24 h, ER stress inducers or UPR inhibitors were treated in the co-culture. Approximately 48 h after the addition of the compounds, cells were subjected to flow cytometric DNA content analysis. For blocking caspase activities, 20 μM zVAD-fmk was administrated to cell cultures 1 h before the administration of ER stressors. zVAD-fmk was newly added every 24h. The two mixed cell populations were separately counted based on the EGFP fluorescence signal.

#### Immunoblotting

Cells were lysed with RIPA buffer (50 mM Tris-HCl, pH 7.6, 150 mM NaCl, 10 mM NaF, 10 mM β-glycerophosphate, 1% NP-40, 0.5% sodium deoxycholate and 0.1% sodium dodecyl sulfate (SDS), protein inhibitor cocktail (cOmplete, Roche)) for 10-15 min on ice then clarified by centrifugation for 10 min with 15,000 rpm. The clarified lysate was mixed with 4 or 5 × SDS-PAGE sample buffer, boiled for 5 or 10 min, and subjected to SDS-PAGE. Separated proteins were transferred onto an Immun-Blot PVDF membrane (Bio-Rad). The blotted membranes were blocked with 0.3% skim milk in TTBS (50 mM Tris, 138 mM NaCl, 2.7 mM KCl, and 0.1% Tween 20), incubated with primary antibodies overnight at 4°C, and incubated with horseradish peroxidase-conjugated secondary antibodies for 2 h at 25°C or overnight at 4°C. Each step was followed by 3 washes with TTBS. For IRE-1 (in Fig.3) and CHOP (in Fig.7) detection, antibodies were diluted with CanGet Signal Immunoreaction Enhancer Solution (Toyobo). The ezWestLumi plus ECL Substrate (ATTO) and a LuminoGraph II chemiluminescent imaging system (ATTO) were used for signal detection.

#### Immunofluorescence and microscopic observations

Cells were fixed with 3.7% paraformaldehyde (Merk) in phosphate-buffered saline (PBS) at room temperature for 10 min, followed by permeabilization with ice-cold 0.5% triton-X100 in PBS containing 0.1 M glycine (Wako) for 5 min on ice. Fixed cells were treated with PBS containing 3% FBS (Gibco) and 3% bovine serum albumin (Wako) for 1 h on ice, incubated with primary antibodies >24 h at 4°C, and with fluorescence-conjugated secondaries >24 h at 4°C at indicated dilutions in PBS containing 5% FBS. DNA was stained with 0.5 μg/mL DAPI (Dojindo). Following each treatment, cells were washed 3 times with PBS.

For detecting protein aggregations, cells were fixed with 4% paraformaldehyde in PBS at 25°C for 30 min, followed by permeabilization with PBS containing 0.5% triton-X100 and 3 mM EDTA (pH 8.0) on ice with gentle shaking for 30 min, and incubated with 1 × dual detection reagent in the Proteostat aggresome detection kit (containing Proteostat aggresome detection reagent and Hoechst 33342; Enzo Life Sciences) at 25°C for 30 min. Following each treatment, cells were washed 1 or 2 times with PBS.

The stained cells were observed under a TE2000 microscope (Nikon) equipped with a ×60 1.4 NA Plan-Apochromatic, a CSU-X1 confocal unit (Yokogawa), and an iXon3 electron multiplier-charge coupled device (EMCCD) camera (Andor) or ORCA-ER CCD camera (Hamamatsu Photonics), or with a Ti2 microscope (Nikon) with ×60 1.4 NA Apochromatic and Zyla4.2 sCMOS camera (Andor). Image acquisition was controlled by µManager (Open Imaging).

#### Colorimetric cell proliferation assay

For cell viability assay, cells were seeded on 96-well plates at 9,000 cells/well. After 24 h, cells were treated with different concentrations of tunicamycin. Forty-four h after the addition of tunicamycin, 5% Cell Counting Kit-8 (Dojindo) was added to the culture, incubated for 4 h, and absorbance at 450 nm was measured using the Sunrise plate reader (Tecan).

#### Statistical analysis

To compare two groups of categorical data or continuous numerical data not assumed to have normal distributions, we used the Brunner-Munzel test. To compare more than two groups of categorical data, we used the Steel-Dwass test. To compare more than two groups of continuous numerical data, we used the ANOVA followed by the post-hoc Tukey test. Statistical significance was set at *p* < 0.05 for all analyses. All statistical analyses were conducted with R software (4.2.1).

## Data availability

The data supporting this study’s findings are available from the corresponding author upon reasonable request.

## Supporting information

This article contains supporting information, including three supplementary figures and one supplementary table.

## Acknowledgment

We thank Fuyu Sato, Ryoto Nomura, and Daiki Saito for their technical assistance.

## Author Contributions

Conceptualization, S.I., K.Ya., E.M., and R.U.; Methodology, S.I., K.Ya., K.S., G.B., E.M., and R.U.; Investigation, S.I., K.Ya, K.S. K.Yo., K.V., E.M., and R.U.; Formal Analysis, S.I., K.Ya., K.S., K.Yo., S.M., G.Y., E.M., and R.U.; Resources, G.B., E.M., and R.U.; Writing – Original Draft, K.Ya., R.U.; Writing – Review & Editing, S.I., K.Ya., and R.U.; Funding Acquisition, S.I., K.Ya., G.B., E.M., and R.U.

## Funding and additional information

This work was supported by JSPS KAKENHI (Grant Numbers 23K19360 and 22KK0110 to Sumire Ishida-Ishihara, JP19J12210 and JP21K20737 to Kan Yaguchi, and JP19KK0181, JP19H05413, JP19H03219, JP21K19244, and JP24K02017 to Ryota Uehara), the Princess Takamatsu Cancer Research Fund, the Orange Foundation, the Smoking Research Foundation, Daiichi Sankyo Foundation of Life Science, the Akiyama Life Science Foundation, and the Terumo Life Science Foundation to Ryota Uehara, and Bilateral Joint Research Projects of Japan Society for the Promotion of Science and Hungarian Academy of Sciences (JPJSBP120193801) to Gabor Banhegyi, Eva Margittai, and Ryota Uehara.

## Conflict of interest

The authors declare that they have no conflicts of interest with the contents of this article.

## Figure legends

**Supplementary figure 1.**
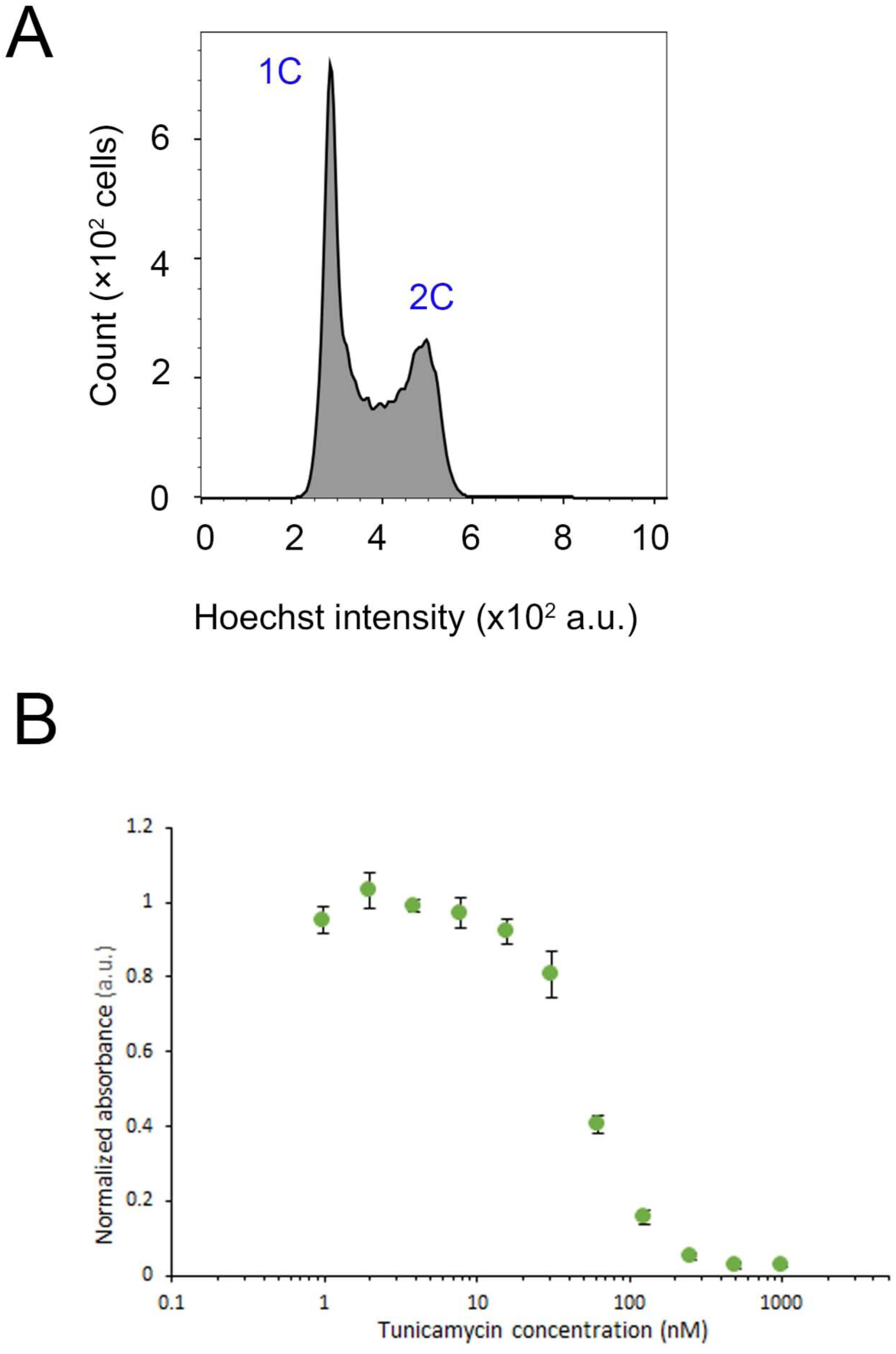
(A) Flow cytometric analysis of DNA content in the haploid cell culture at 20 h after thawing. Representative data from 2 independent experiments. **(B)** Dose-response curves of normalized absorbance in a colorimetric cell proliferation assay in haploid HAP1 cells treated with tunicamycin. Mean ± SE of 6 replicates from 3 independent experiments.

**Supplementary figure 2.**
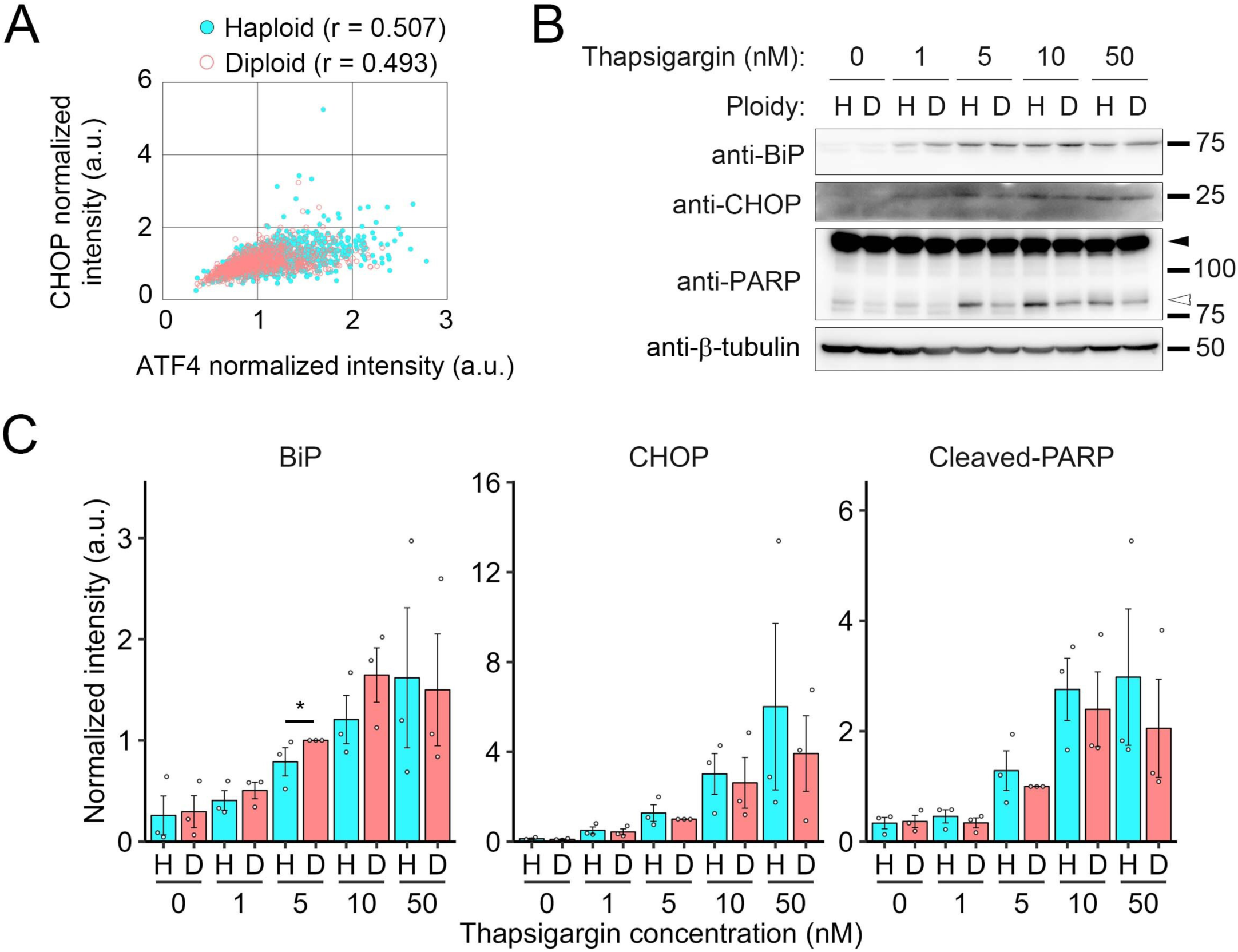
(A) A dot plot of CHOP signal against ATF4 signal in the immunostaining of haploids and diploids treated with tunicamycin for 24 h with correlation coefficient. Data are identical to those in Fig. 3D. **(A)** Immunoblotting of BiP, CHOP, and PARP in haploid or diploid HAP1 cells treated with different concentrations of thapsigargin for 24 h. The closed or open arrowhead indicates uncleaved or cleaved PARP, respectively. β-tubulin was detected as a loading control. Representative results from 3 independent experiments. **(C)** Quantification of the relative amount of BiP, CHOP, or cleaved PARP. Mean ± SE of 3 independent experiments. Asterisks indicate statistically significant differences among samples (\**p* < 0.05, the Brunner-Munzel test).

**Supplementary figure 3.**
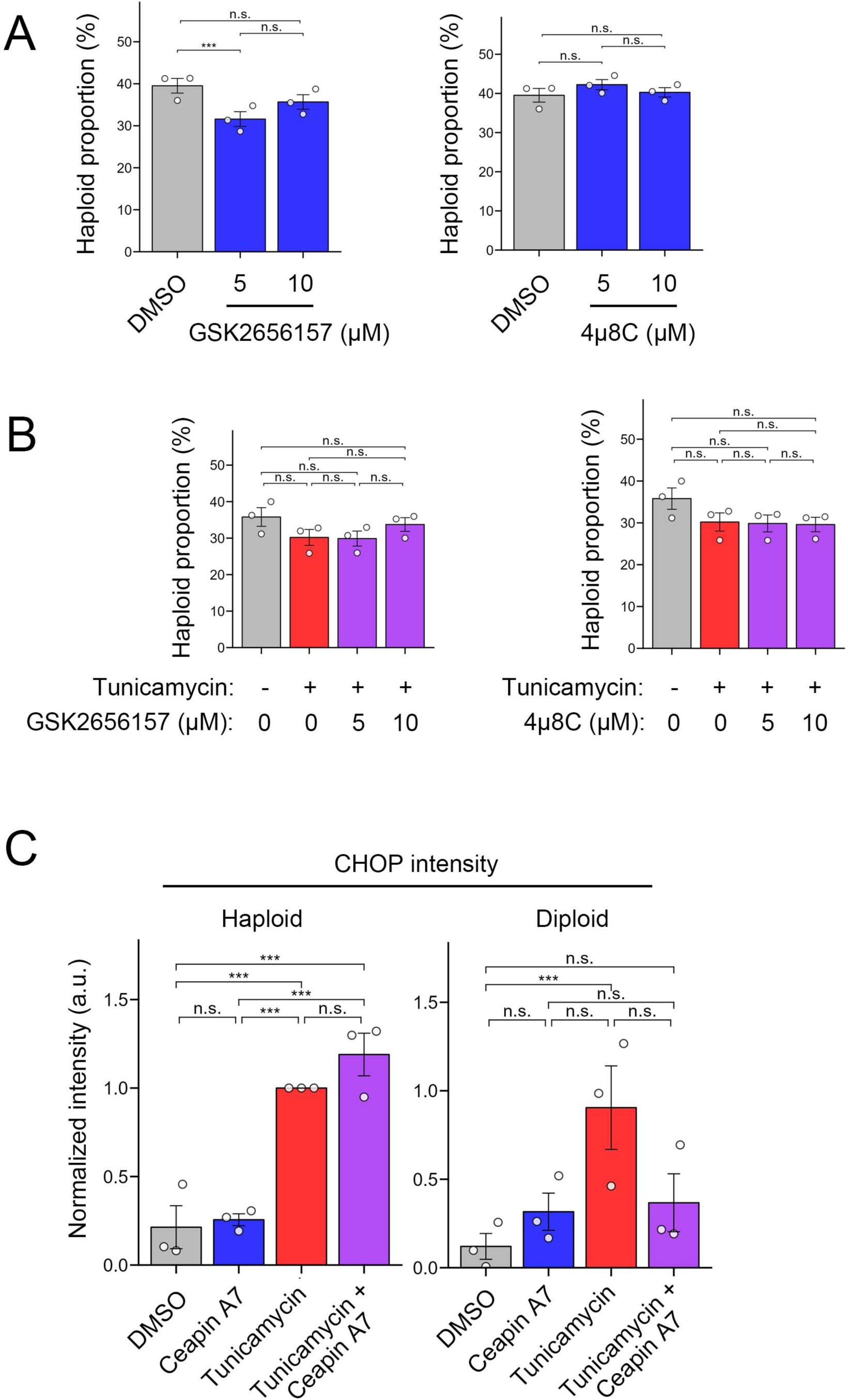
(A, B) The proportion of haploid cells in the haploid-diploid co-culture treated with different UPR inhibitors (A) or co-treated with UPR inhibitors and tunicamycin (B). Mean ± SE of 3 independent experiments for each condition. Asterisks indicate statistically significant differences between conditions (\*\*\**p* < 0.001, the Steel-Dwass test). For comparison, the same control data (DMSO in A, or DMSO and tunicamycin in B) as in Fig. 7A or B are shown in each graph in A or B, respectively. **(A)** Quantification of the relative intensity of CHOP in the immunoblotting analysis in Fig. 7C. Mean ± SE of 3 independent experiments. Asterisks indicate statistically significant differences among samples (\*\*\**p* < 0.001, the Steel-Dwass test).

## Notes

### Competing Interest Statement

The authors have declared no competing interest.

