## Supplementary material for "Fragility of ER homeostatic regulation underlies haploid instability in human somatic cells": Table S1

Table S1: A list of antibodies used in this study

| **Host and antigen** | **Identifier and distributor** | **Dilution** | **Application** |
| --- | --- | --- | --- |
| Rabbit monoclonal anti-ATF4 | D48B, Cell signaling Technology | 1:1000  1:500 | IB  IF |
| Mouse monoclonal anti-ATF6 | ab122897, abcam | 1:1000 | IB |
| Rabbit monoclonal anti-BiP | C50B12, Cell Signaling Technology | 1:1000 | IB |
| Mouse monoclonal anti-CHOP | L63F7, Cell Signaling Technology | 1:1000  1:500 | IB  IF |
| Rabbit polyclonal anti-Grp94 | #2104, Cell Signaling Technology | 1:1000 | IB |
| Rabbit polyclonal anti-PARP | #9542, Cell Signaling Technology | 1:1000  1:5000 | IB  IB (Fig.7) |
| Rabbit monoclonal anti-PERK | #3192, Cell Signaling Technology | 1:1000 | IB |
| Rabbit monoclonal anti-IRE1 | #3294, Cell Signaling Technology | 1:1000 | IB |
| Mouse monoclonal anti-β-tubulin | 10G10, Wako | 1:1000  1:10000 | IB  IB (Fig.7) |
| Goat peroxidase affinipure anti-rabbit IgG (H+L) | 111-035-003, Jackson Immuno Research Laboratory Inc. | 1:1000  1:5000 | IB  IB (Fig.7, PARP) |
| Goat peroxidase affinipure anti-mouse IgG (H+L) | 115-035-003, Jackson Immuno Research Laboratory Inc. | 1:1000  1:5000  1:10000 | IB  IB (Fig. 7, CHOP)  IB (Fig. 7, β-tubulin) |
